# A Framework to Determine Active Connectivity within the Mouse Brain

**DOI:** 10.1101/2023.12.27.573396

**Authors:** Guanhua Sun, Tomoyuki Mano, Shoi Shi, Alvin Li, Koji Ode, Alex Rosi-Andersen, Steven A Brown, Hiroki Ueda, Konstantinos Kompotis, Daniel Forger

## Abstract

Tremendous effort has focused on determining the physical connectivity within the mouse brain. However, the strength of connections within the brain constantly changes throughout the 24-hour day. Here, we combine experimental and computational methods to determine an “active connectivity” of the physical connections between the most active neurons. Brain cells of freely behaving mice are genetically marked with the activity-dependent TRAP2 system, imaged, digitized, and their connectivity is inferred from the latest brain atlases. We apply our methods to determine the most active networks in the early light and early dark hours of the day, two periods with distinct differences in sleep, wake, and feeding behavior. Increased signaling is seen through the visceral and agranular insular (AI) regions in the early day as peripheral stimuli are integrated. On the other hand, there is an increase in the activity of the retrosplenial cortex (RSP) and the anterior cingulate cortex (ACC) during the early night, when more sustained attention is required. Our framework carves a window to the three-dimensional networks of active connections in the mouse brain that underlie spontaneous behaviors or responses to environmental changes, thus providing the basis for direct computer simulations and analysis of such networks in the future.

## Introduction

Brain activity throughout the 24-hour day is sculpted by the functional interactions of neuronal ensembles[1]. Understanding these complex behaviors involves exploring which sets of neurons are “on duty” within a given temporal window and how they interact. Much effort has focused on mapping out the physical connectivity of the mouse brain with both experimental and statistical methods[2–5]. However, physical connectivity does not necessarily determine the dynamics of brain functions. Brain organization has been reported to vary in a time-dependent manner in both animal and human studies, from the level of molecules in the individual cells to functional circuits [6–8]. As a result, the networks that decide brain-wide behaviors, for instance, at different times of the day, emerge as a complex result of the physical connectivity itself and whether the connected counterparts are active or quiescent. In the human brain, core neurocognitive networks, such as the salience, default mode, and central executive network (SN, DMN, and CEN), have been identified chiefly by employing resting-state functional magnetic resonance imaging (fMRI). Interestingly, the interaction of these networks has been demonstrated to regulate attention and interoception, as well as mental fatigue, in the rodent brain, which consists of homologous regions, as in primates.

At the same time, most brain functions involve more than one brain area or specific kind of neuronal circuit. For example, the sleep-wake cycle affects the activity of almost all major brain regions: there are wake-promoting circuits from the thalamus to the cortex and circuits from the midbrain to the cortex or thalamus that promote sleep[9]. In the context of sleep regulation, a multi-structural “state-switch” has been well established[10]. Nevertheless, how these structures communicate with each other and other cortical or subcortical areas at different times of the day has thus far been understudied. Although those behaviors are generated by neuronal circuits across different regions, only specific sub-groups of neurons in different regions are often involved [11, 12]. Identifying the neuronal populations responsible for specific behaviors within those regions remains challenging.

To tackle this limitation, neuronal tagging systems based on genes transcribed shortly after neuronal activation, namely immediate early genes[13], have been developed[14, 15]. More specifically, the Targeted Recombination in Active Populations (TRAP), or the TRAP2 system, utilizes transcription factor(TF) binding under the *Fos* promoter to express an improved Cre (iCre) recombinase enzyme that is active only in the presence of the synthetic drug 4-hydroxytamoxifen (4-OHT). Combining the TRAP2 system with a Cre-dependent fluorescent marker enables labeling neurons active at a specific temporal window or behavior [16, 17] several hours after injection. Subsequently, novel tissue-clearing technologies, such as CUBIC[18], combined with light sheet microscopy, allow researchers to visualize and map the most active neurons within the three-dimensional (3D) brain.

But how can such discrete neuronal data be further used? We propose a method to reconstruct the active neurons and map their connections digitally. Several current tools can help us quantify and reconstruct 3D brain structures with individual neuron positions mapped into a common brain coordinate framework, such as the Allen Brain Atlas Common Coordinate Framework (ABA-CCF) [19] or the CUBIC atlas [20]. Recently, investigators have shown that the whole mouse brain can be scanned and the position of each neuron’s cell body determined[20]. Here, we apply a similar machine-learning

approach to identify TRAP2-captured neurons in the mouse brain and map them to the ABA. With neurons’ anatomical information aligned with the ABA-CCF, we can use available connectomes associated with the atlas to build connectivity. A mesoscale connectome of the mouse brain from thousands of injection experiments is available [2]. This work was extended to a statistical model to calculate the brain connectivity matrix to the resolution of a voxel that is 100\mu m^˄^3, which is the finest mouse whole-brain connectome to date [3]. We use this voxelized connectome model to determine the probability of connections between our identified active neurons. We also integrate another cell atlas’s excitatory/inhibitory information [21]. Finally, we combine all information and apply a probabilistic algorithm for regions of interest to complete the connectivity, which is immediately visualized and can be simulated by different computational platforms [22] [23] [24] [25] or analyzed [26].

To validate this framework, we apply our method to four separate brains, two where neurons were “TRAPed” during the early hours of daylight (Zeitgeber Time 0; ZT0), and two during the hours after the onset of darkness (ZT12), each marking active neurons over the next 4 hours. We then determine the positions of the cell bodies in each brain using a machine-learning method and map them to the ABA. After the 3D identification of the positions of the cell bodies of neurons, we analyze the most active regions at two different time points and construct connectivity related to those regions, where our results are aligned with various experimental findings. We also build a whole-cortex connectivity using the same algorithm for computational simulations.

## Results

### Modular framework for connectivity construction

In the current work, we present a modular framework to quickly build detailed inter-regional connectivity between the active neuronal circuits during any chosen time period or behavior. The framework is divided into three modules:

#### TRAP2 system to tag Fos expressing neurons

First, we need experimental techniques to tag the neurons or circuits of interest genetically. Here, we focus on labeling active neurons at different time points of the day. We consider mice during the beginning of the day (ZT0) and the beginning of the night (ZT12). The differences in connectivity between these two time points could be caused by the changes in light level, sleep needs, hunger, and circadian phase (**Fig 1a**).

**Fig 1:**
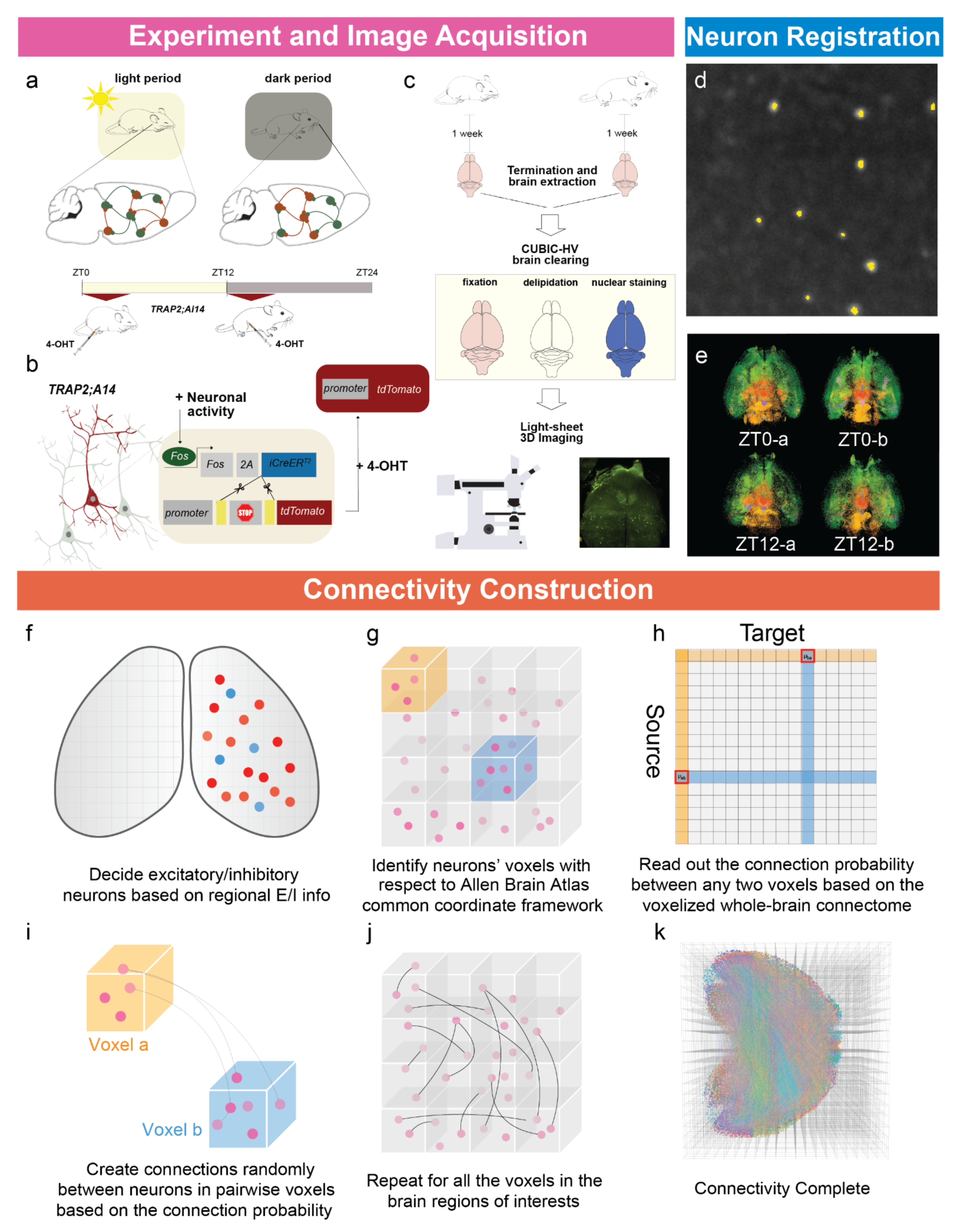
Workflow of our experimental and computational framework. **a:** Four TRAP2 - tdTomato expressing mice are prepared under the 12:12 Light-Dark schedule, where we determine which neurons are most active at two timepoints. **b:** We administered 4-OHT to two mice at ZT-0, subsequently collecting the subjects 4 hours later at ZT4. Simultaneously, the remaining two mice were administered at ZT12 and collected at ZT16. Administration of 4-OHT to TRAP2- tdTomato mice is able to activate Cre recombinase to cleave the stop cassette, thus allowing for tdTomato expression to show neuron’s activity level under different behaviors. **c:** Brains are extracted after a week for sufficient tdTomato expression and undergo the CUBIC-HV clearing method before light-sheet microscopy and 3D image acquisition. **d:** From the light-sheet images, neuron bodies are segmented using machine-learning techniques. Neurons’ physical positions are identified and aligned to the ABA- CCFv3. **e:** In the end, we obtain four whole-brain datasets with neurons identified as points. **Top**: two brains where 4-OHT were administered at ZT0(ZT0-a/b). **Bottom**: two brains where 4-OHT were administered at ZT12(ZT12-a/b). **f-k:** Connectivity between neurons is constructed with the following steps: **(f)** Excitatory and inhibitory neurons are identified based on the regional E/I information. **(g-h)** Based on the voxelized connectome, the neuron’s position is used to determine its probability of connecting with other neurons. **(i-k)** The voxel-based algorithm is repeated for regions of interest until the connectivity is completed.

Our TRAP system, where c-Fos is used as the marker for neuron activities, tags neurons active at ZT0 or ZT12. Four TRAP2 (aka *Fos^2A-iCreERT2^* ; see **Methods**) mice, which were bred with Ai14 (aka Ai14^(RCL-tdT)-D^; in-house breeding) mice, are prepared under 12-12 light-dark conditions. 4-Hydroxytamoxifen were administered to two mice either at ZT0 or ZT12, and allow expression of c-Fos through Z T-0 to ZT4 or ZT12 to ZT16. Brains are then collected at ZT4 or ZT16 and stored for seven days to allow sufficient tdTomato expression (**Fig 1b**). After that, the brains are cleared following CUBIC protocols, and brain images are acquired using plane illumination microscopy techniques (**Fig 1c**).

#### Segmentation and reconstruction

With neuronal images properly stitched, we now apply machine learning techniques to identify neuron bodies and their positions. We use a two-step convolutional cell detection algorithm implemented for the CUBIC Atlas[20] to identify both the position and the strength of the c-Fos expression level (see **Methods**). After identifying all neurons, we align our data to the ABA-CCFv3 to use the incorporated connectivity information (**Figs 1d-e**).

#### Connectivity construction

The Allen Brain Atlas has multiple levels of refinement. It has divided the brain into voxels of different sizes, and the voxelized model interpolates the strength of connectivity between any voxels with size of 100\mu m^˄^3 based on the previous mesoscale connectome[2]. We will utilize this voxelized connectivity matrix [3] to connect neuron populations located within the regions of our interest.

We also need to determine if a neuron is excitatory or inhibitory to produce a more complete connectome for it to be simulated. As ABA does not give excitatory and inhibitory information, we also use another cell atlas [21], which provides the number of excitatory and inhibitory neurons for all major brain regions. See the method section for more details.

Now, we propose a novel algorithm that utilizes the aforementioned information to build the connectivity for *N* neurons in the regions of our interest. Let 𝑋_!_ represent the physical positions of the ith neuron and W the voxelized connectivity matrix, where 𝑊_"#_ represents the connection probability from voxel 𝑎 to voxel 𝑏. Our connectivity algorithm has one parameter 𝜌 : the average number of connections per neuron. With this information, we create the connectome as follows (**Figs 1f-k**):

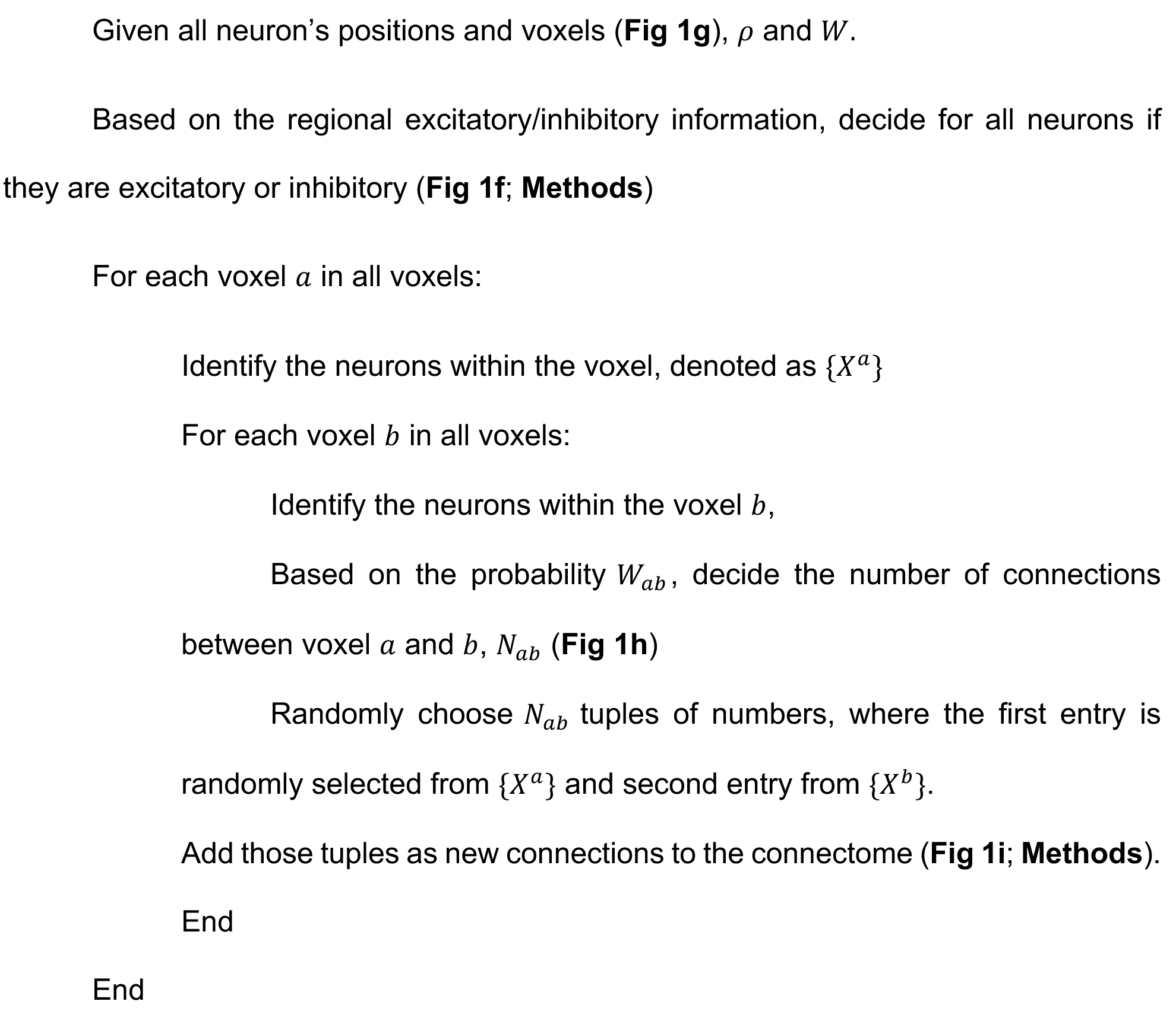

The algorithm is repeated for all pairwise voxels in our interest regions until the connectivity is completed (**Figs 1j-k**). Since the computation time can significantly increase if there are many neurons, two computational methods here are used to substantially speed up the process of generating connectivity and storing them. When developing the connectivity, unlike most algorithms that iterate through the neurons, we loop through the spatial voxels for a significant speed up. As a result, the above-proposed algorithm can be two orders of magnitude faster than iterating through the neurons naively (see **Methods**). Secondly, sparse matrix representations are used to store the generated connectivity, exploiting that the fact that realistic connectivity is almost always sparse even if the number of neurons is large. Such representations of connectivity are also compatible with other computational packages such as PyGeNN[22] and NEURON[23].

### Neuronal information of the brains at different time points

To demonstrate the usefulness of our pipeline, we consider two time points with opposing wake and sleep behavior, after light and after dark onset (injections at ZT0 and ZT12, respectively). These are studied in mice where we genetically marked all active (as indicated by c-fos expression) over the next 4 hours. We first visualized all the neuronal data we collected in this process. We look at the 3D reconstruction of all four brains (**Fig S1**) with neurons identified as single points with their positions and regional information aligned with the ABA-CCFv3. We visualize the first brain collected at ZT0(ZT0a) as an example in **Figs 2a-c**. We also look into the difference between the regional composition of the four brains, where the number of neurons of each region (**Fig 2d**) and its corresponding percentage (**Fig 2e**) is calculated. We immediately see a time-point difference between the four brains: both morning brains have more c-fos expressed neurons than evening (**Fig 2d**) across the whole brain. This greater number of active neurons could indicate a higher level of fatigue due to the accumulation of activities during the night. On the other hand, the percentage of cortical neurons to the total number of neurons in the evening brains is larger than in the morning.

**Fig 2:**
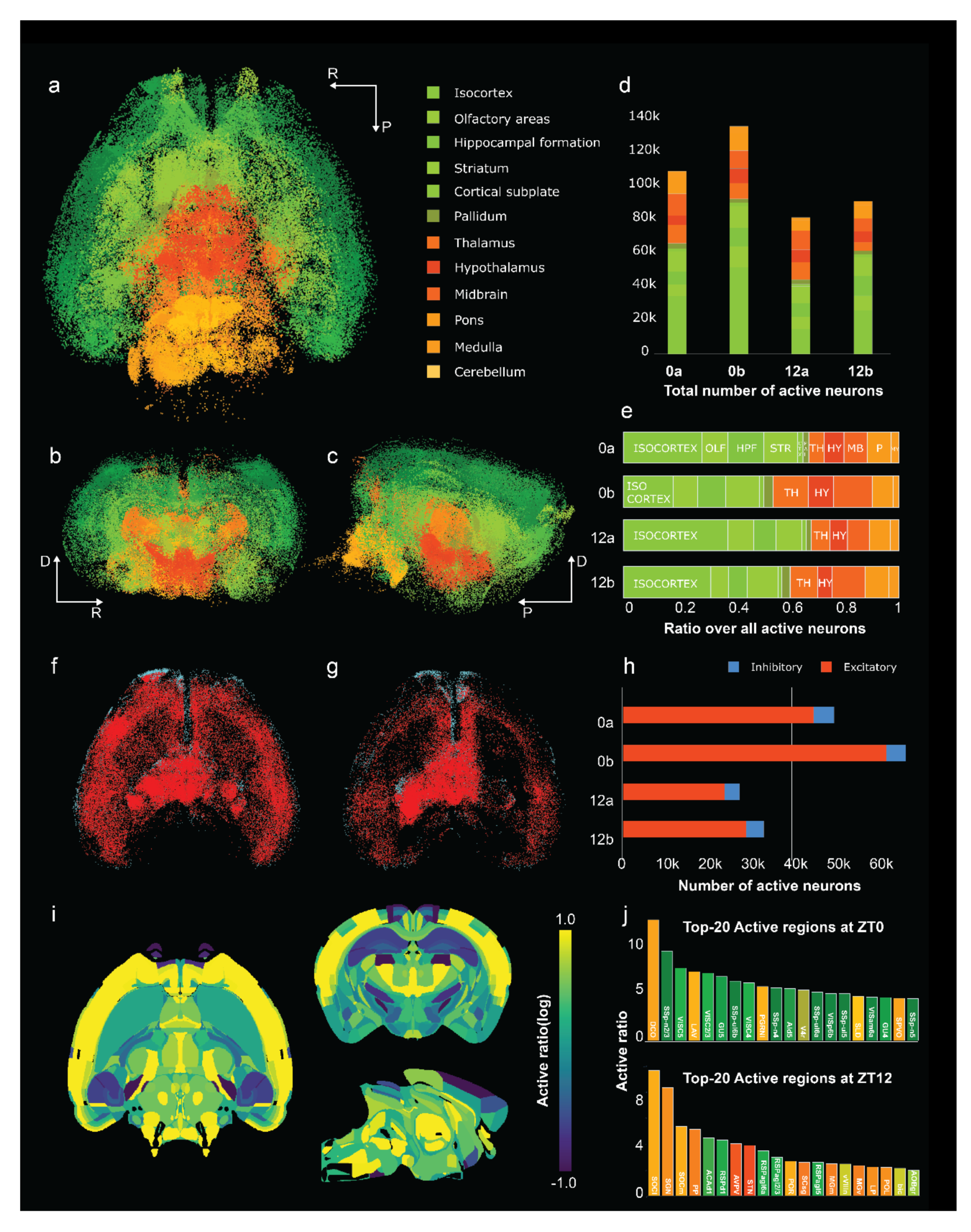
Whole-brain neuronal information of 4 brains collected at 2 different time points. **a-c:** The horizontal, coronal and sagittal view of the neurons identified in the first brain collected at ZT0(ZT0-a). The colors are determined by the Allen Brain Atlas colormap based on the regional information. Arrows indicate the directions of anterior(A)- posterior(P), dorsal(D)-ventral(V) and left(L)-right(R). **d:** The number of neurons identified in all four brains: ZT0a, ZT0b, ZT12a and ZT12b. While there are some individual differences, the number of active neurons identified in both brains at ZT0 is larger than those collected at ZT12. **e:** The percentage of neurons in each of the different brain regions. **f-g:** The isocortex, thalamus and hypothalamus of two brains(**f: ZT0a, g: ZT12a**) visualized based on the excitatory/inhibitory information. **h**: The number of excitatory and inhibitory neurons identified in the isocortex, thalamus, and hypothalamus at four different time points. **i:** Horizontal, coronal, and sagittal slices at the center of the whole brain, colored based on the index ratios that show whether the regions are active at ZT-0 or ZT-12. **j**: The top20 active regions at ZT0(**Top**) and ZT12(**Bottom**).

From the regional information provided by a cell atlas[21], we can classify neurons as excitatory and inhibitory across the cortex, thalamus, and hypothalamus based on our algorithm previously described. The results of two brains after light onset and two brains after light offset are shown in **Figs 2f-g**. Though our excitatory/inhibitory information is static for each region, the number of neurons identified in each region differs between the two time points. Therefore, each brain’s ratio of excitatory to inhibitory neurons is different (**Fig 2h**). In this figure, we observe that the number of excitatory neurons changes drastically compared to the number of active inhibitory neurons, which are relatively unchanged.

Next, we locate the most active or inactive regions at the two time points for further analysis. Since we have two brains at each time point, we first calculate the average number of active neurons for each region at one time point with data from two brains. We then calculate the ratio of the average number of neurons of each region in the morning over the ratio in the evening. The larger this ratio, the more active the region is after light onset to after light offset. We color the brains based on this index ratio (**Fig 2i**) and present the 20 most active regions after light onset (ZT0) and after light offset (ZT12) in **Fig 2j**. Tost active regions after light onset are mostly cortical, in particular visceral, gustatory, and agranular insular regions. On the other hand, the most active regions after light offset are mostly subcortical, except large parts of the retrosplenial cortex and some layers of the anterior cingulate cortex.

### Active visceral and agranular insular connectivity is higher after lights on

As nocturnal animals, mice are spontaneously less active during the light period. After light onset (ZT0), the animals have been mainly awake during the night and tend to minimize locomotor activity and feeding. While our study did not include sleep deprivation, we note that many of the regions that were most active either at ZT0 or ZT12 in **Fig 2j**, including the AI, ACC, and RSP, also were found to have higher *Fos* activation in a study [27] looking at increased *Fos* activation due to sleep deprivation, suggesting that the changes we are seeing are related due to cognitive fatigue.

A common phenomenon with increased cognitive fatigue is a lesser tolerance for pain and an increased sensitivity to signaling from the periphery. As cognitive fatigue increases, we become more sensitive to hunger, pain, other signaling, and emotional regulation[28]. Thus, we hypothesized that input to the brain would be increased after lights were on, which would be processed via the thalamic-insula pathway. Indeed, some of the largest changes in connections involve this pathway in our data. **Fig 3a** shows increased signaling from the agranular insular cortex to other brain parts. The two brains from the light-on timepoint showed upregulation of this pathway over the two brains at lights off. For simplicity, we show one representative brain for each time point and condition going forward. The input to the AI from other brain parts was also greatly increased after the lights were on (**Fig 3b**). We also found this increased signaling to and from the visceral cortex, likely due to signals linked to feeding behavior.

**Fig 3:**
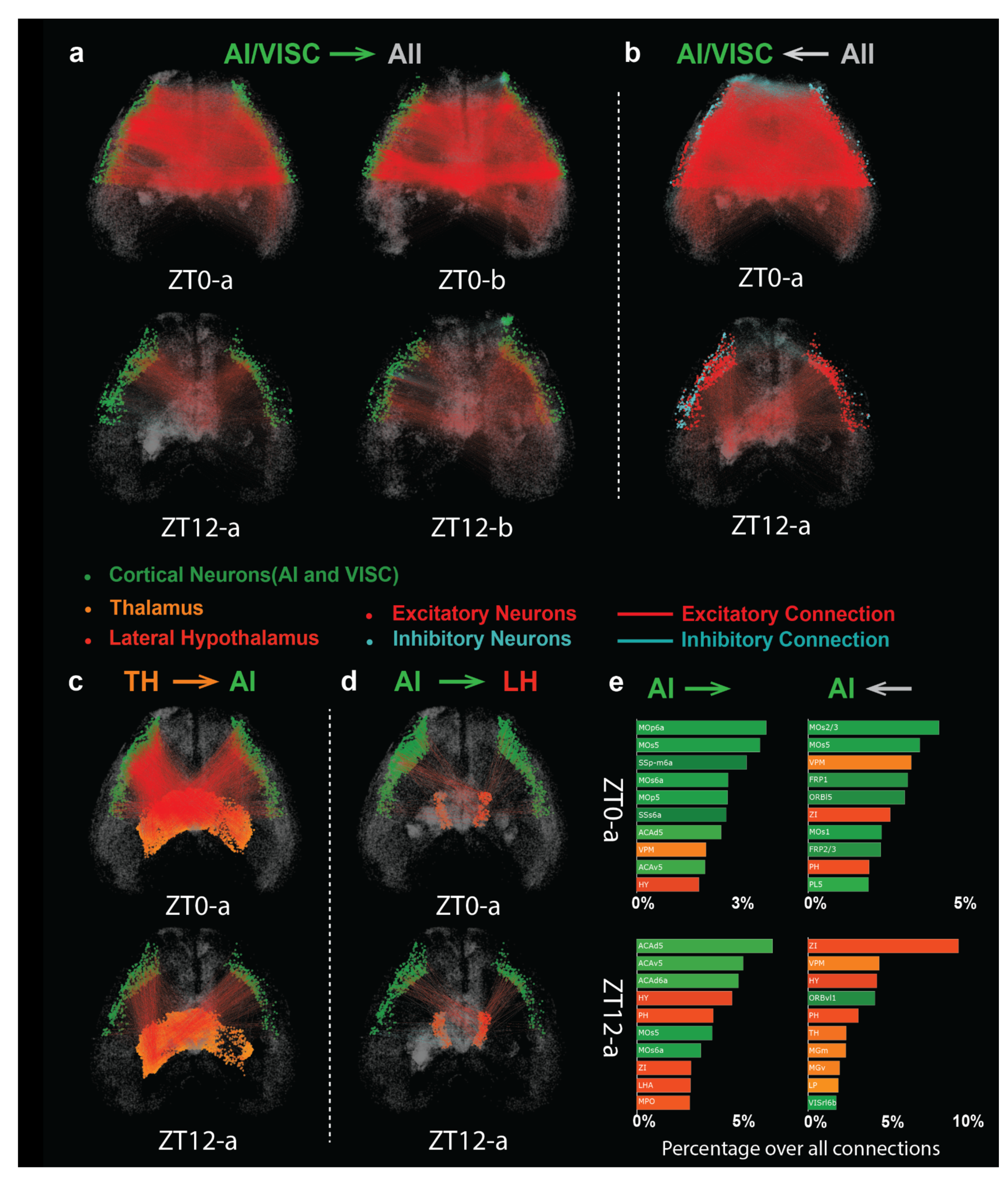
Upregulation of visceral and agranular insular connectivity in the early day. **a:** Projections from agranular insular cortex (AI) and visceral regions (VISC) to the whole brain. **Top:** ZT0-a/b. **Bottom:** ZT12-a/b. A sharp contrast is visible from the visualizations. **b:** Projections from the whole brain to AI and VISC in two brains(**Top:** ZT0a, **Bottom:** ZT12a), where neurons are colored differently here to show their excitatory/inhibitory information. **c:** Projections from the thalamus (TH) to AI. (**Top:** ZT-0a, **Bottom:** ZT-12a) **d**: Projections from AI to the lateral hypothalamus (LHA)(**Top:** ZT-0a, **Bottom:** ZT-12a). **e:** Bar plots that show the top regions that AI projects to **(Left)** or project to AI **(Right)** at different time points (**Top:** ZT-0a, **Bottom:** ZT-12a).

We next tried to look for a specific pathway within this circuit. Recent work has highlighted signaling from the thalamus to the insula and the lateral hypothalamus in regulating behaviors related to seeking food [29]. We explored these pathways in **Fig 3c**, which shows signaling from the thalamus to the AI increases after lights on, likely due to the overall increased signaling due to cognitive fatigue. Indeed, it is well known that thalamic signaling increases with sleep deprivation, although overall activation of the insula (rather than specific to the AI) may decrease [30]. However, the actual food seeking behaviors would align with the ZT12 -active neurons as feeding occurs during lights-on. We see that the signaling from the AI to the lateral hypothalamus (LH) is similar across the lights-on and lights-off cases (**Fig 3d**).

In contrast, most other regions have a markedly decreased signaling at ZT12 from the AI. This could be because of increased activity in the LH due to food entrainable circadian oscillators. Overall, we find that while signaling to and from the AI is decreased after lights off, this decrease is much less in signaling between thalamus and hypothalamus and more from cortical areas (**Fig 3e**). Thus, this network balances increased hunger signaling and cognitive fatigue after lights on, with increased circadian activation in the LH after lights off and signaling from other cortical areas.

### Active retrosplenial connectivity in the early evening indicates higher attention needs/default mode network activation

Unlike most cortical regions that are less active after lights off, some subregions of the retrosplenial cortex(RSP) and the anterior cingulate cortex(ACC) show much higher activity after lights off. Both regions are thought to be related to the default mode network (DMN) [31], and activation of the ACC is important in sustained attention [32]. Since ZT12- ZT16 marks the beginning of the night, we hypothesized that there would be more connections between the ACC and RSP, which shows a higher level activation of the DMN [33]. Indeed, as shown in **Fig 4a**, connections between the ACC and the RSP are more numerous after the lights are off rather than after the lights are on.

**Fig 4:**
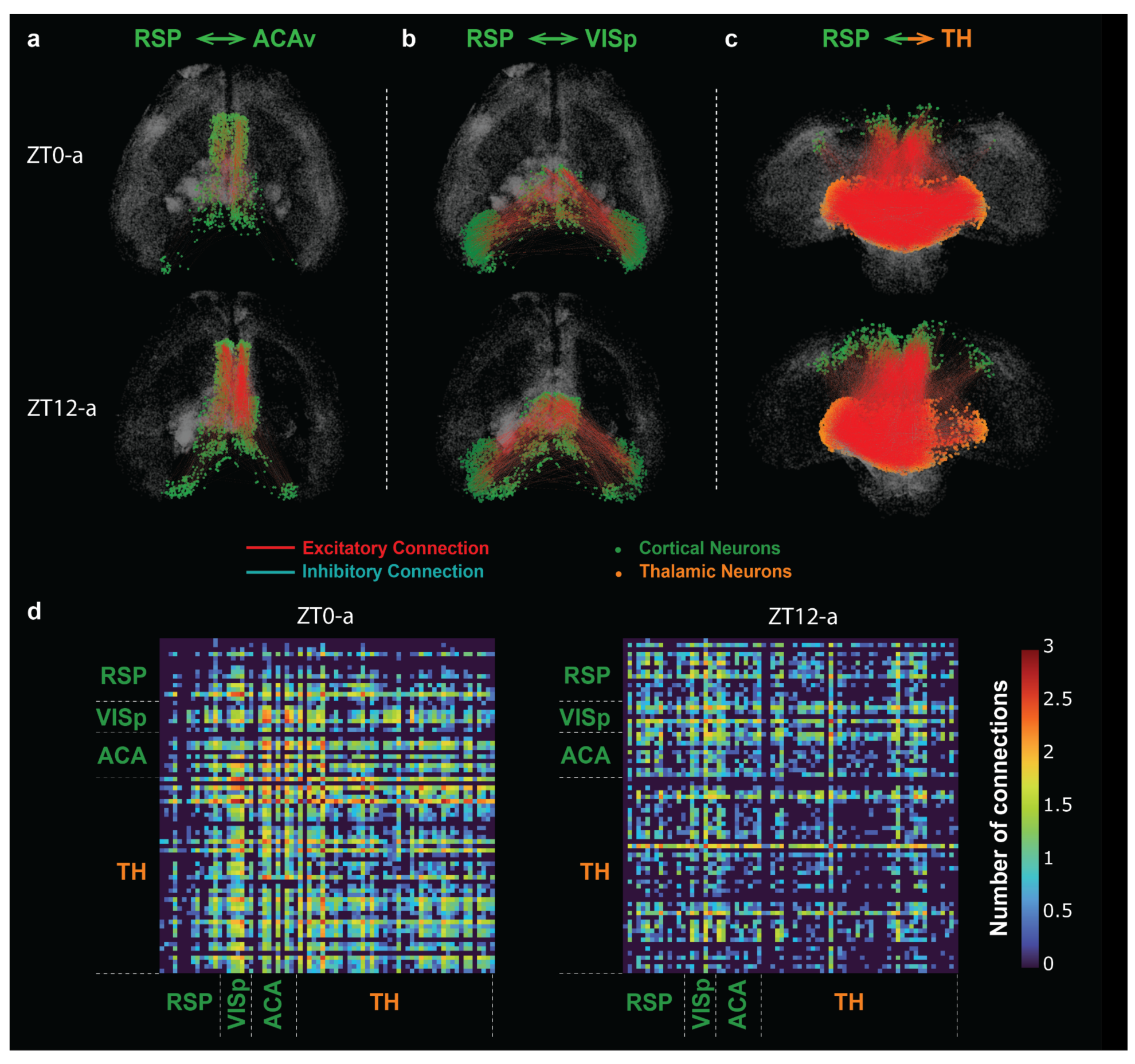
Upregulation of retrosplenial connectivity in the early evening. **a:** Connections between the retrosplenial cortex (RSP) and the anterior cingulate cortex (ACC) (**Top:** ZT0-a, **Bottom:** ZT12-a). Here, the connectivity is visually more active at ZT12 than the opposite situation presented in Fig3A. **b:** Connections between the RSP and the primary visual cortex (VISp) (**Top:** ZT0-a, **Bottom:** ZT12-a) Here, the connectivity is on the same level because the active neurons in RSP are up-regulated at ZT12 but the neurons in VSIp are down-regulated at ZT12. **c:** Connections between the RSP and the thalamus (TH). (**Top:** ZT0a, **Bottom:** ZT12a) **d:** The connectivity matrix that shows the number of connections between RSP, VSIp, ACA, and TH in logarithmic scale (**Left:** ZT0a, **Right:** ZT12a).

For comparison, in **Fig 4b**, we study connections between the RSP and the primary visual cortex (VISp). Neurons in VISp are upregulated in the morning because the lights are on and downregulated after the lights are off. This is balanced against the increase in RSP signaling after lights off, so the overall number of active connections is relatively similar between the two conditions. We also see similar numbers of active connections between the RSP and the thalamus after lights on and after lights off (**Fig 4c**) again since there is greater thalamic activation after ZT0 due to cognitive fatigue but greater RSP activation after ZT12. **Fig 4d** plots more details of these connections. These connections could be part of the spatial navigation pathways that must work throughout the day [34–36].

We also note some asymmetries between the left and right connections that we found (e.g., **Fig 4c, bottom**). Similar left-right asymmetry has also been observed in fMRI studies of fatigue or in the signaling of visceral stimuli to the anterior insula to control feeding behavior [37]. Further work could study the asymmetries in terms of cognitive fatigue.

## Discussion

The current work is the first to analyze functional brain connectivity from an activity-dependent genetic tagging of neurons with brain clearing and 3D imaging, integrating E-I neuronal information. We create a modular, openly accessible pipeline in freely behaving mice to determine networks that are responsible for behaviors. As a proof of principle, we study structures active at the beginning hours of light or darkness. Notably, the brain regions we find most changed at different times of day align with a recent study on Fos expression in the context of a chronic sleep deprivation paradigm [27]. Based on these results, our pipeline should be able to determine the neuronal networks responsible for many additional behaviors.

While nocturnal animals are more fatigued in the early morning than in early evening, there are many other physiological processes that change over the course of a day. Light-directed responses or circadian circuit components are also represented in our dataset. To the best of our knowledge, our most active structures during daytime have not been associated with the direct effects of light in literature. Conversely, we report that the most active structures during the “daytime” (ZT0-4) in our study are mostly linked to interoception [38], which has been suggested to increase with, as well as affect, sleep behavior[39]. More specifically, the insula has been characterized as a hub where large-scale brain networks converge[40]. It occupies an established, pivotal role in Default Mode Network (DMN), as well as Salience Network (SN) in the brain[41, 42]. In the context of sleep, weaker DMN connectivity has been linked to poorer sleep in human adolescents[43]. Reduced AI activation and functional connectivity in humans have been linked to less sleep amounts and more disadvantageous choices during risk-taking. In contrast, sleep duration was positively correlated with increased functional coupling (FC) between the AI and dorsolateral prefrontal cortex under high-stress risk-taking[44] . Lesions in the agranular cortex of the insula have been found to regulate sleep and wake distribution[45]. Its importance for physiology, cognitive processing, and sleep has been further underlined by its simultaneous increased vulnerability to a-synucleinopathy and its potential contribution to the deterioration of sleep patterns in Parkinsonian patients[46]. Conversely, dark-phase active structures in our study were more associated with locomotor behavior, spatial signal integration, and cognitive alertness. The RSP is crucial during spatial navigation, particularly in the darkness[47]. More specifically, it involves path integration[35, 36] and the formation of navigational memories. As such, it communicates with other space-relevant structures, such as the hippocampus and anterior thalamic nucleus.

Elduayen and colleagues[47] have shown that the RSP cortex is necessary for path integration and incorporating visuospatial information, particularly during darkness. Mis-timed increased connectivity between the retrosplenial cortex and the hippocampus leads to objective sleep disturbances in patients suffering from psychiatric disorders due to pathophysiological increased DMN connectivity[48]. Furthermore, a rodent study integrating molecular and anatomical signatures of sleep and wake behavior found Homer1a and Jun upregulated in the retrosplenial cortex only after spontaneous or forced wakefulness, compared to sleeping controls[49].

At the same time, the ACC has been causally implicated in motivation and decision-making [50], while its stimulation has been reported to inhibit fear response[51]. In the context of sleep, increased ACC connectivity has been associated with awakenings during N2 insomnias[52], and has thus been deemed an area of high interest for primary insomnia[53]. In the study from Thompson and colleagues, [49], neuronal activation marker FosL2 was found to be increased in ACC after both spontaneous wakefulness in the dark and forced wakefulness in the first 6 hours of light period. The third most activated structure in our dark period dataset was the visual cortex, which is necessary for spatial navigation. Indeed, V1 activity is regulated by locomotion, distance head movements and gain[54], suggesting a potential increase in connectivity with the anterior thalamic area, which is highly implicated in spatial navigation and gain control.

Several limitations of our methodology should be noted. First, the cell-finding algorithms may not be perfect. For this reason, we compare the relative ratio of neurons between the two time points to decide which regions are most active rather than the absolute number. We also do not consider the relative amount of *c-Fos* activation, although that could provide additional information in future studies. That being said, when the algorithm is trained on multiple brains, it yields similar numbers of neurons across all brains, which suggests we are capturing the vast majority of *c-Fos*-positive cells. The machine learning algorithms can be tuned to determine as many neurons in a particular brain. In supplemental data, we provide one example of ∼400,000 neurons detected for brain ZT0-a(**Fig S2**). However, they opted for a stricter learning paradigm, which gives ∼100,000 neurons detected in each brain but provides an algorithm that consistently works across all brains studied at once.

Secondly, while the statistical model of the physical brain connectivity represents a heroic effort, it does not represent a complete mapping of the neurons within the mouse brain. Even if these could be measured, recording the billions of connections within the brain presents data storage issues. The voxelized connectivity model Knox et al. provided is the finest mouse connectome, but it relies on data with much coarse resolution[2]. It becomes tricky when we try to use the finer interpolated connectivity matrix: if we look at the hypothalamus connectivity matrix provided by the computational model, we do see that the connectivity matrix is very striped, i.e., many post-synaptic neurons receive very homogeneous input from all up-stream neurons. This is not ideal since we are not getting the inhomogeneity we wish to get from the voxelized model. At the same time, this connectivity does not map out local connectivity or microcircuitry within each voxel. As such connectivity is relatively unknown, special experiments are required, or statistical models can also be used to infer them [5]. Additionally, such microcircuitry can be later added to our model later. Better connectivity data might be needed to build finer local connectivity. Thus, building local connections with currently available data remains challenging. However, with many researchers pursuing this, new connectivity data will soon become available, which could easily be incorporated into our methods to refine connectivity matrices. Finally, we assume that the physical connections between neurons, as indicated by the Allen Brain Atlas, do not change over the day but rather the strength of these physical connections. While the strength of synapses is much more variable than the physical connectivity, we can not rule out possible changes to physical connectivity. These could be explored with future connectivity work. Again, our method is agnostic to the Atlas or connectivity data used.

As networks responsible for behaviors are identified with our pipeline, the neuronal basis of the physiology of these behaviors can be explored. Active connectivities could also be visualized, analyzed, and used as the basis of detailed neuronal simulations. A package to do this is described in a companion to this manuscript. Neuronal data of the four brains are also provided for further analysis and could be used in platforms such as NEURON[23], BRIAN[24], NEST[55], and packages such as PyGeNN[22]. Thus, our pipeline should provide a starting point for many future investigations.

## Methods

### Animal preparation

For this experiment, *TRAP2* (aka Fos^2A-iCreERT2^ ; JAX #030323) mice were bred with *Ai14* (aka Ai14^(RCL-tdT)-D^; inhouse breeding) mice. Male 10-14 weeks old mice, heterozygous for both alleles, were used for experiments according to the Experimental Design below (also see **Figs 1a-c**). Animals were housed according to relevant regulations from the veterinary office of Canton Zurich. All mice used were males, housed in groups in polycarbonate cages (31 × 18 × 18 cm) in a sound attenuated and temperature/humidity-controlled room (23-24°C, 50-60% respectively). Mice were kept under a 12h light/ 12h dark cycle (light intensity 70-90 lux) and had access to food and water *ad libitum*. Genotyping for the Fos2A-iCreER alleles was performed according to published protocols[17]. All experiments were approved by the animal welfare officers of the University of Zurich and the relevant veterinary authorities of the Canton of Zurich, Switzerland (license national number 32308 and 34563).

### 4-OHT preparation and administration

For 4- Hydroxytamoxifen preparation we followed a protocol developed by the Deisseroth lab (https://web.stanford.edu/group/dlab/optogenetics/sequence_info.html). Briefly, 4-OHT powder was diluted in 250ul DMSO (Sigma H6278). Subsequently, the diluted 4-OHT was mixed with 4.75 ml of separately prepared 25% Tween80 diluted in saline, and mixed by vigorous vortexing.

The resulting clear solution was used within the next 1 hour for intraperitoneal (i.p) injections of 250 μl volume per animal to deliver a final concentration of 20 mg/kg 4-OHT.

### Experimental design

Tagging of active neurons at ZT0-4 or ZT12-16 was achieved via i.p. injection with 20 mg/kg 4- OHT in saline at the onset of the respective periods (**Fig 1b**). After allowing seven days[17] for the expression of tdTomato in captured neurons, mice were terminated with the pentobarbital administration and perfused at the middle of the initial trapping window (ZT2 and ZT14, respectively). Brains were extracted preserving all structures, and underwent the CUBIC tissue-clearing process.

### CUBIC brain clearing

Briefly, whole brains were immersed in 4% paraformaldehyde (PFA) for 24h, then delipidated and decolored for 5 days in CUBIC-L (CUBIC-HV^TM^1, TCI chemicals). Subsequently, their cellular nuclei were stained with DAPI for another 5 days, and the refracting index (RI; 1.520) of the transparent brain was calibrated in CUBIC-R+ solution (CUBIC-HV^TM^1, TCI chemicals). Following the end of incubations and in-between washes, the cleared brains were stored for two days in mineral oil (Mounting Solution (RI 1.520), TCI chemicals) and imaged in the following days.

### Whole-brain imaging

The mesoscale selective plane illumination microscopy instrument (mesoSPIM, https://mesospim.org/; [56] was used for axially-scanned light-sheet imaging of the cleared brain.

Transparent whole brain images were recorded at 3.2x magnification with an Olympus MVX-10 macroscope with a MVPLAPO objective combined with a Hamamatsu ORCA-FLASH 4.0 V3 camera, at a voxel size of 2.03 × 2.03 × 5 μm3 (X × Y × Z). The laser/filter combinations for imaging were as follows: for registration of autofluorescence and light-scattering, a 405nm (100mW) excitation laser at 50% and no emission filter. For the tdTomato signal, a 561nm (100mW) excitation laser at 50% and a quadruple bandpass (BP444/27; BP523/22; BP594/20; BP704/46) emission filter. Brains were positioned such that the ventral portion was the first plane acquired. Briefly, each whole brain was imaged in 16 tiles per channel (eight tiles per illumination; right or left). The acquired data ranged around 400GB per brain. Subsequently, images were stitched and fused separately for each channel using the FIJI plugin BigStitcher [57], according to the “*MesoSPIM stitching using BigStitcher*” guidelines (https://zmb.dozuki.com/Guide/MesoSPIM+Stitching+using+BigStitcher/283). Finally, whole brain images were stored in .tiff format for further processing in the CUBIC pipeline.

### Cell detection

To detect and quantify neurons labeled with tdTomato from 3D brain images acquired with light-sheet microscopy, we used the cell detection method described in [58. First, the raw TIFF image was converted into HDF5 format to allow fast and parallel access to the 3D array data on disk. Then, a supervised machine learning method implemented in ilastik{Berg, 2019 #394] was used to classify voxels into three categories: (1) tdTomato-expressing cells (2) fibers labelled with fluorescent proteins or blood vessels with strong autofluorescence and (3) background. Following this classification, single neurons were segmented and fluorescence intensities were quantified using ecc (https://github.com/DSPsleeporg/ecc).

### Brain registration

To register cleared brain images to the atlas space, we followed and implemented the method used in the CUBIC pipeline [59] [20] [58]. We used Allen Mouse Brain Common Coordinate Framework (ABA-CCF) v3 as a reference brain [19]. We downsampled the autofluorescence channel of the cleared brain to make a reduced image with 50 um the voxel size. We then aligned it with the average brain template image of CCFv3 (50 um voxel size) using the symmetric image normalization method (SyN) implemented in ANTs library [60]. Since portions of the olfactory bulb (OLF) and cerebellum (CB) were obstructed in our image, we excluded these regions from our analysis. To do this, we manually made image masks to hide OLF and CB from the cleared tissue image. Accordingly, a modified version of ABA-CCFv3 was generated using a custom Python script where OLF and CB were hidden.

After computing the warp filed to map cleared brain to the atlas, we applied the same transformations to the table of the detected cells. We then assigned a unique brain region ID to each cell using the annotation image provided by ABA-CCFv3.

### Use of excitatory/inhibitory information

E/I ratios of major brain regions are provided in [21] but they are in a slightly different format and resolution compared to ABA-CCFv3. For each neuron, if the regional E/I ratio is not given, we go one level-up until the E/I ratio is provided by the atlas. After obtaining the E/I ratio of the region that the neuron belongs to, we randomly decide whether the neuron is excitatory or inhibitory. For example, if the E/I ratio of a region is r, then we say the probability for a neuron in this region to be excitatory will be 𝑝_!"#_ = r/(r+1), and 𝑝_$%&_ = 1/(r+1). In this actual algorithm, say a region has N neurons, we will randomly assign N*𝑝_!"#_ neurons to be excitatory and the rest are inhibitory. All neurons are initiated to be excitatory or inhibitory before building the connectivity. but is of coarser resolution than the voxelized positional data so we have to coarse grain to utilize the cell atlas.

### Normalizing the connectivity matrix and choose **𝜌**

The connectivity matrix provided by the voxelized connectivity model was normalized according to the values of the whole brain and therefore the values are not to be used directly to build regional connections between the neurons. We develop a single parameter algorithm which allows us to use the values in the connectivity matrix as a probability or distribution to determine the connectivity. Assume we have identified N neurons in M voxels, where the connectivity matrix W is M*2M, including both the contralateral and ipsilateral connections. We first calculate the average of the connectivity matrix W, denoted as 𝜌. If we assume each voxel has the same number of neurons and each neuron has the same probability 𝜌 to connect with other neurons, then the average number of connections per neuron will be 𝑁 ∗ 𝜌. Then we have to decide the average number of connections we want for each neuron and let’s denote this number to be 𝜌. To get the number we want, we now multiply every entry of W by 𝜌/𝜌 such that now the average number of connections per neuron will be \rho. Since the number of neurons in each voxel is different and they have different probabilities to connect with each other, the result is somehow inaccurate but will be on the same level of magnitude.

### Details of the connectivity algorithm

In the connectivity algorithm, the crucial step is to determine the number of connections between voxels and choose appropriate up and down-stream targets. Consider we are building a connectivity for N neurons, which posit in M voxels that span different regions. Naively, we can iterate through each neuron and read out the probability to connect with other neurons based on the voxel that contains the neuron. This algorithm will take O(N*N) time, which is very expensive for a large number of neurons. Here, we optimize the algorithm by iterating through the voxels that these neurons sit in. Amazingly, this proposed algorithm will take O(ρM*M) time, where M is the number of voxels that contain all the neurons, and this number has a maximum, based on the voxel resolution we choose. If we assume the number of neurons in each voxel is roughly the same, In the end this algorithm is about two orders of magnitude faster than iterating through the neurons.

## Data visualization

The connectivity is visualized with Python package Plotly. All colormaps, if not specially noted, follow the colormap provided by the Allen Brain Atlas CCFv3.

## Acknowledgments

We thank Ningyuan Wang for helpful discussions on this work.

We acknowledge the following funding: HFSP RGP0019/2018 to DBF, SB and HU, NSF DMS 2052499 to DBF, and ARO MURI W911NF-22-1-0223 to DBF

---

**Fig S1:**
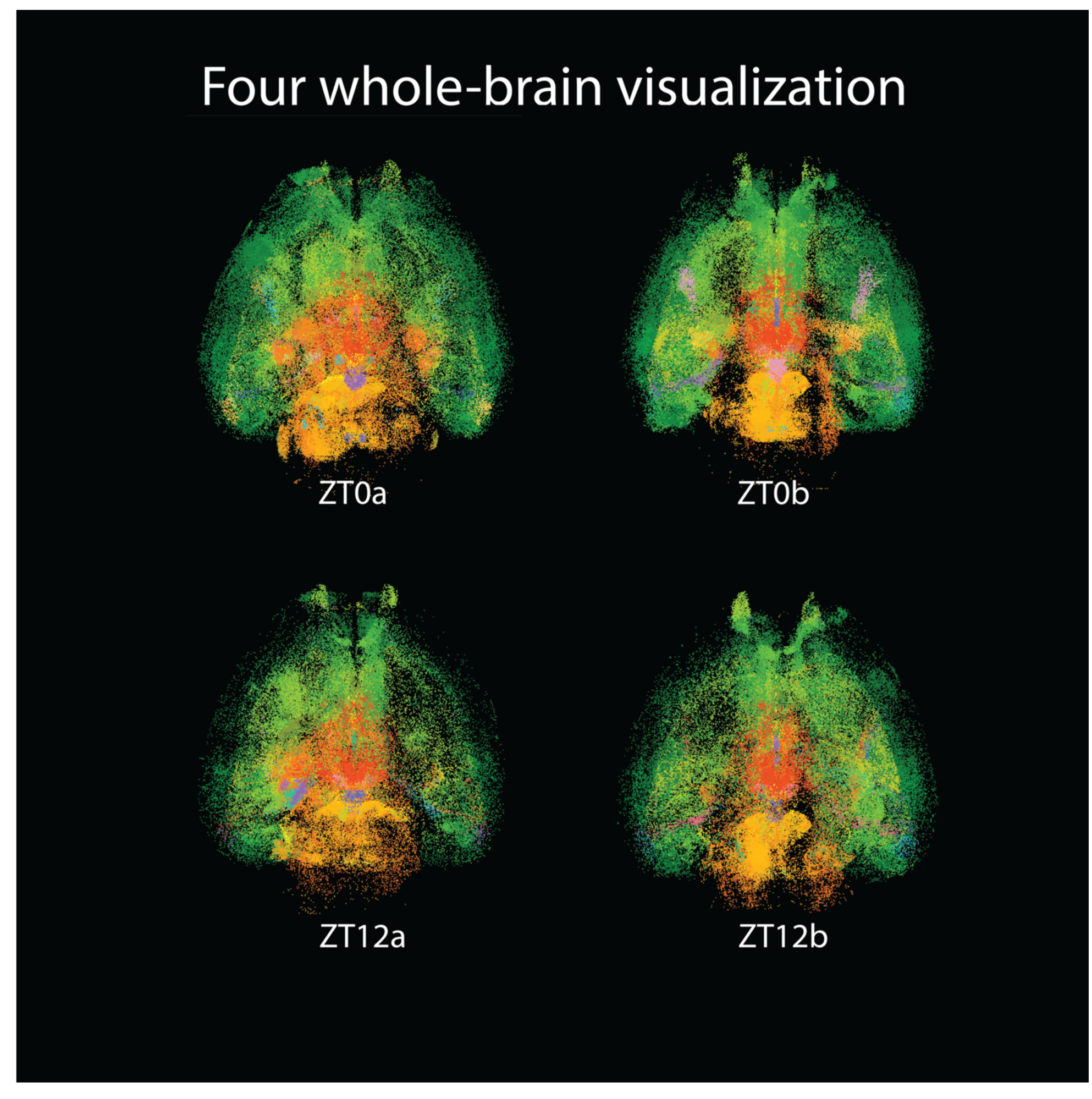
Whole-brain visualization of the four brains we collected. **Top: ZT0a/b Bottom: ZT12a/b**

**Fig S2:**
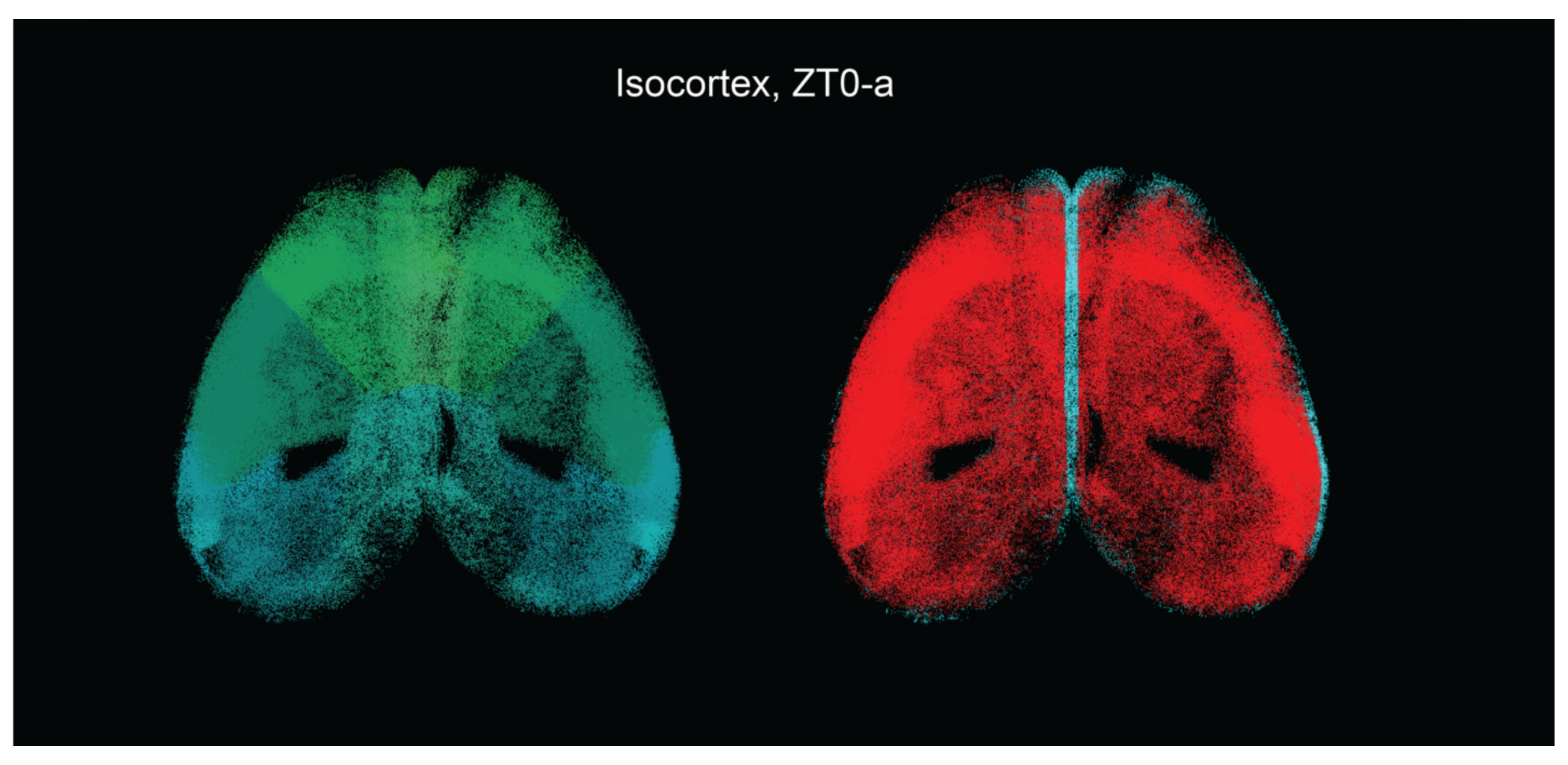
Visualization of the isocortex of ZT0-a, where an alternative algorithm is tailored for this brain and more than 400,000 neurons are detected. **Left:** Colored based on regional information. **Right:** Colored based on E/I information.

